# T-cell distribution in the dorsal root ganglion across species, sex, and age

**DOI:** 10.64898/2026.04.30.722027

**Authors:** Jayden A. O’Brien, Nishka Kuttanna, Khadijah Mazhar, Marisol Mancilla Moreno, Asta Arendt-Tranholm, Joseph B. Lesnak, Michael A. Wilde, Spoorthi Sadasivuni, Pooja J. Patel, Rainer V. Haberberger, Armen N. Akopian, Stephanie Hennen, Verena Arndt, Jane M. Brandon, Katherin A. Gabriel, Seph M. Palomino, Amol M. Patwardhan, Theodore J. Price

## Abstract

T-cells infiltrate somatosensory ganglia in response to nerve damage, autoimmune disease, and infection, contributing to sensory abnormalities and pain. In naïve states, T-cells are rare in the rodent dorsal root ganglion (DRG) but have been reported in human and non-human primates without known relevant exposures. It remains unclear whether there are inherent evolutionary or species differences in DRG T-cell residence. Using a comparative biology approach, we investigated the frequency and distribution of T-cells in the mammalian DRG across humans, non-human primates, pigs, and rodents, and in humans investigated the contributions of sex and age. Spatial transcriptomics and immunofluorescence independently verified the robust presence of DRG T-cells at similar levels in humans, non-human primates, and pigs, but were fewer in rats and largely absent in mice. In humans, premenopausal females were more likely to have elevated DRG endoneurial T-cells than post-menopausal females or adult males. T-cells were detected in human dorsal root ganglion at as early as two months of age but were less abundant within the perineuronal niche. Most human DRG T-cells expressed distinct markers consistent with a resident memory (Trm) phenotype. We discuss the importance of studying the functional roles of DRG-resident T-cells and raise broader considerations for modelling peripheral nervous system disease.

## Introduction

T-cells are key directors of immunity in homeostasis and disease whose heterogeneity enables targeted antigen-specific responses. Naïve T-cells continuously recirculate between peripheral blood and secondary lymphoid organs. Upon successful T-cell receptor stimulation with cognate antigen, these cells undergo expansion and maturation, gain effector and eventually memory function, and are more likely to infiltrate or circulate through nonlymphoid tissues where they can further recruit other immune cells to direct a targeted immune response. Tissue-resident memory T-cells (Trm), on the other hand, are long-lived cells present in some nonlymphoid tissues that do not typically recirculate and have distinct surface markers, transcriptomes, and functional traits from circulating memory cells [14; 62; 71; 84]. They are most studied at barrier sites, such as the intestines, lung, and skin, where they are phenotypically shaped by the local milieu, ignore egress signals, and are responsible for potent location-specific recall in response to antigen reencounter [62].

Somatosensory ganglia, including the dorsal root ganglia (DRG) and trigeminal ganglia, are peripheral nervous system structures containing the cell bodies of primary sensory neurons responsible for transducing touch, temperature, nociception, and proprioception from cutaneous and visceral tissues. These sensory neurons innervate barrier tissues such as skin and gut and thereby innervate notable nonlymphoid sites of neuroimmune interaction. T-cells that associate with sensory neurons – either at the distal nerve ending, within the nerve, or within the ganglion itself – both release and respond to neuropeptides and other efferent molecules to maintain homeostasis or contribute to pathology [6; 24; 26; 48; 73]. These pathologies include sensory and motor autoimmune neuropathies [58; 69], autoimmune diseases such as multiple sclerosis [83], neuropathic pain [18; 31; 32; 61], and herpes simplex and varicella-zoster viruses which can become latent within somatosensory ganglia [10; 70; 76; 77; 79].

Our understanding of the role of DRG T-cells in human disease and homeostasis relies primarily on mouse and rat models. In naïve rodents, T-cells are generally considered to be negligibly present in somatosensory ganglia [34; 41; 44]. However, T-cell infiltration into the rodent DRG is observed to occur following nerve injury in both mice [30; 32; 80] and rats [4; 5; 27; 28; 30; 32; 55; 63; 80], and the degree of T-cell infiltration into somatosensory ganglia appears dependent both on sex and time elapsed after injury [31]. T-cells also infiltrate the rodent somatosensory ganglia in a wide variety of other conditions, including in the context of chemotherapy exposure [21; 41], infection [43; 57], diabetes [2], and in experimental autoimmune encephalomyelitis [83].

In contrast, human somatosensory ganglia are reported to contain numerous T-cells, even in those without a reported history of peripheral nervous system disease or injury [9; 19; 23; 29; 61; 75]. They have likewise been reported in non-human primates [35]. This discrepancy raises the possibility that there may be inherent species differences in DRG T-cell residency. This has implications for better understanding species-specific contributors to Trm development and for the use of rodent models for modeling human peripheral nervous system disease. Systematic analysis of T-cell density using a consistent quantification approach between species has not previously been conducted to shed light on this question.

Sex differences may also contribute to differences in DRG T-cell density and function, for example due to influences by sex hormones [20; 36; 59; 60]. The presence of T-cells in DRG may also be affected by age, given that the T-cell repertoire shifts from naïve to memory-dominant over time and their abundance and phenotype are associated with aging-associated inflammation in other tissues [20; 64; 67; 85].

We aimed to systematically determine the species-specific distribution of T-cells in the DRG by evaluating T-cell abundance from 82 unique individuals across seven species using spatial transcriptomics, immunofluorescence, flow cytometry, histology, and electron microscopy. T-cell abundances were compared across humans; non-human primates including rhesus macaque, cynomolgus, and baboon; pigs; rats; and mice free of any known autoimmune, inflammatory, or neurological conditions or interventions. We also sought to characterize the baseline frequency, phenotype, and intercellular interactions in human DRG T-cells and whether there is appreciable variation along the dimensions of sex and age.

## Results

### T-cells are robustly present in the human dorsal root ganglion

The experimental workflow is outlined in **Figure 1**. This study began with the observation from previously published single-nucleus RNA sequencing of human DRG data [9] that T-cells form a distinct cell cluster across multiple donors (**Figure 2A**). To verify this finding, we performed fluorescence-activated cell sorting for CD45^+^, CD11b^+^, and CD3^+^ events on enzymatically dissociated cells from 36 perfused human DRGs originating from 20 unique organ donors (**Figure 2B**). Of the CD45^+^ cell fraction, CD3^+^ cells were present in all donors, though there was substantial variability in the absolute frequency of CD11b^+^ and CD3^+^ cells recovered. On average, approximately 100,000 CD11b^+^CD45^+^ cells and 10,000 CD3^+^CD45^+^ cells were recovered per DRG (**Figure 2C)**. Accordingly, CD3^+^ cells represented slightly less than 10% of the overall CD45^+^ cell compartment in the DRG. There were no significant differences in frequency noted between females and males in this measure, though only four female samples were evaluated (**Figure 2D**).

**Figure 1.**
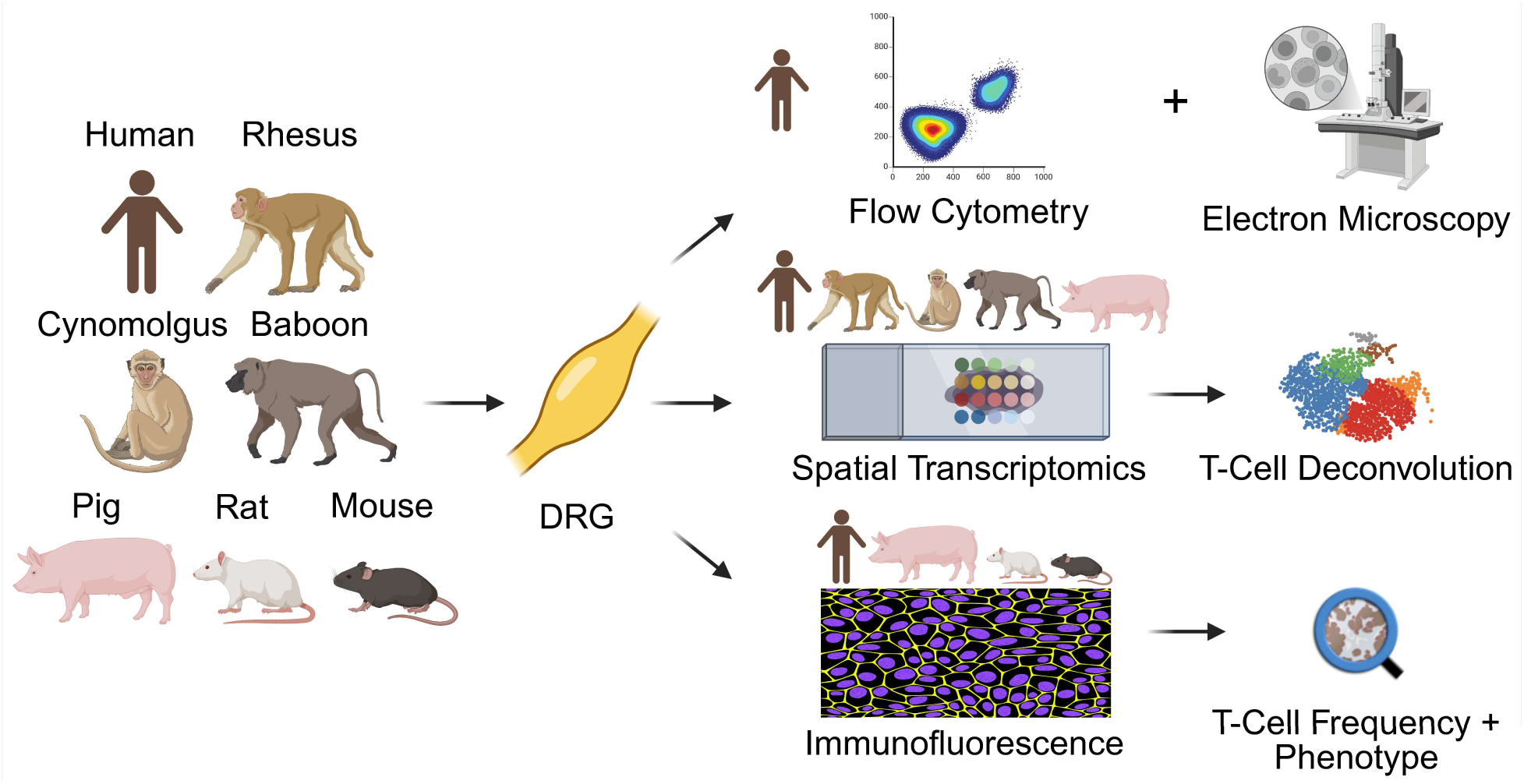
Outline of the experimental workflow and methods used for each included species.

**Figure 2.**
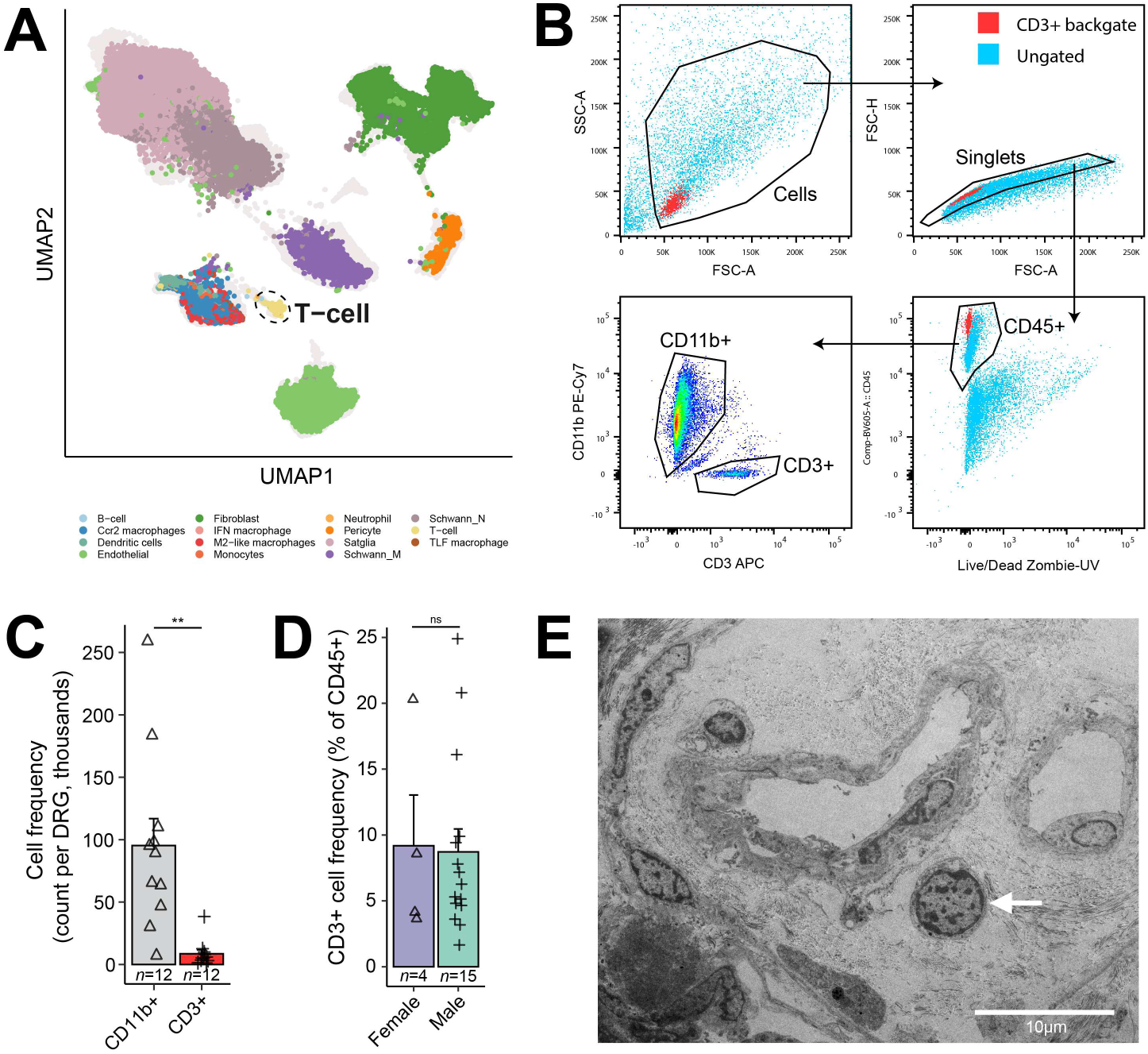
T-cells are present in the human dorsal root ganglia, as detected by single-nucleus RNA sequencing, flow cytometry, and transmission electron microscopy. **(A)** T-cells cluster separately in a previously published single-nucleus RNA sequencing atlas of human dorsal root ganglion non-neuronal cells [9]. Generated at https://painseq.shinyapps.io/harmonized_painseq_v2. **(B)** Fluorescence-activated cell sorting gating strategy from a representative donor. The CD11b-CD3 gate is pseudocolored by cell density. Other plots are colored according to CD3^+^ backgating. **(C)** The cell frequency of CD11b^+^CD45^+^ cells (mean: 92115; range: 7145–259065) and CD3^+^CD45^+^ cells (mean: 8591; range: 1149–38413) isolated from human DRG. CD3^+^ T-cells were approximately 10-fold less abundant than CD11b^+^ myeloid cells (Welch’s unequal variance *t*-test, *t_11.5_* = 4.34, *P* = 0.0011). **(D)** The frequency of isolated CD3^+^ T-cells as a percentage of all live CD45^+^ DRG cells. No differences were noted between sexes (Welch’s unequal variance *t*-test, *t_4.3_* = 0.11, *P* = 0.92). **(E)** Electron photomicrograph showing a lymphocyte in the perivascular space surrounding two profiles of small vessels. \*\**P* < 0.01, ns: not significant. Numbers below columns denote group *n*.

Since these T-cells could be present within the DRG epineurium or other extraneous tissues included for digestion, we next used transmission electron microscopy to verify the presence of lymphocytes within the intact DRG parenchyma. This analysis identified lymphocytes within DRG tissue, often amid small perivascular spaces. Defining features of these lymphocytes included an approximate 5 µm diameter, a round morphology, and the presence of prominent heterochromatin (**Figure 2E**). These results confirm the presence of T-cells in the human DRG across multiple individual donors.

### T-cell transcriptomic signatures are distributed through the DRG of large mammals

We next sought to more systematically investigate the *in situ* abundance and spatial distribution of T-cells within the DRG and compare across species using existing spatial transcriptomic data sets (Visium V1). Given resolution limitations on small tissue sections, we restricted analysis to large mammals including human (*Homo sapiens*); non-human primate including rhesus macaque (*Macaca mulatta*), cynomolgus monkey (*Macaca fascicularis*), and olive baboon (*Papio anubis*); and pig (*Sus scrofa*). Initial inspection of human DRG spatial transcriptomics data used raw counts of 10 T-cell-related genes from a manually curated list (**Supplemental Table 1**) to determine barcode positivity for T-cells.

We found that barcodes positive for this T-cell gene signature were dispersed throughout the DRG, with a small proportion of barcodes containing a strong T-cell signature that colocalized with paraneuronal nucleus-dense regions in the corresponding histological stain (**Figure 3A**).

**Figure 3.**
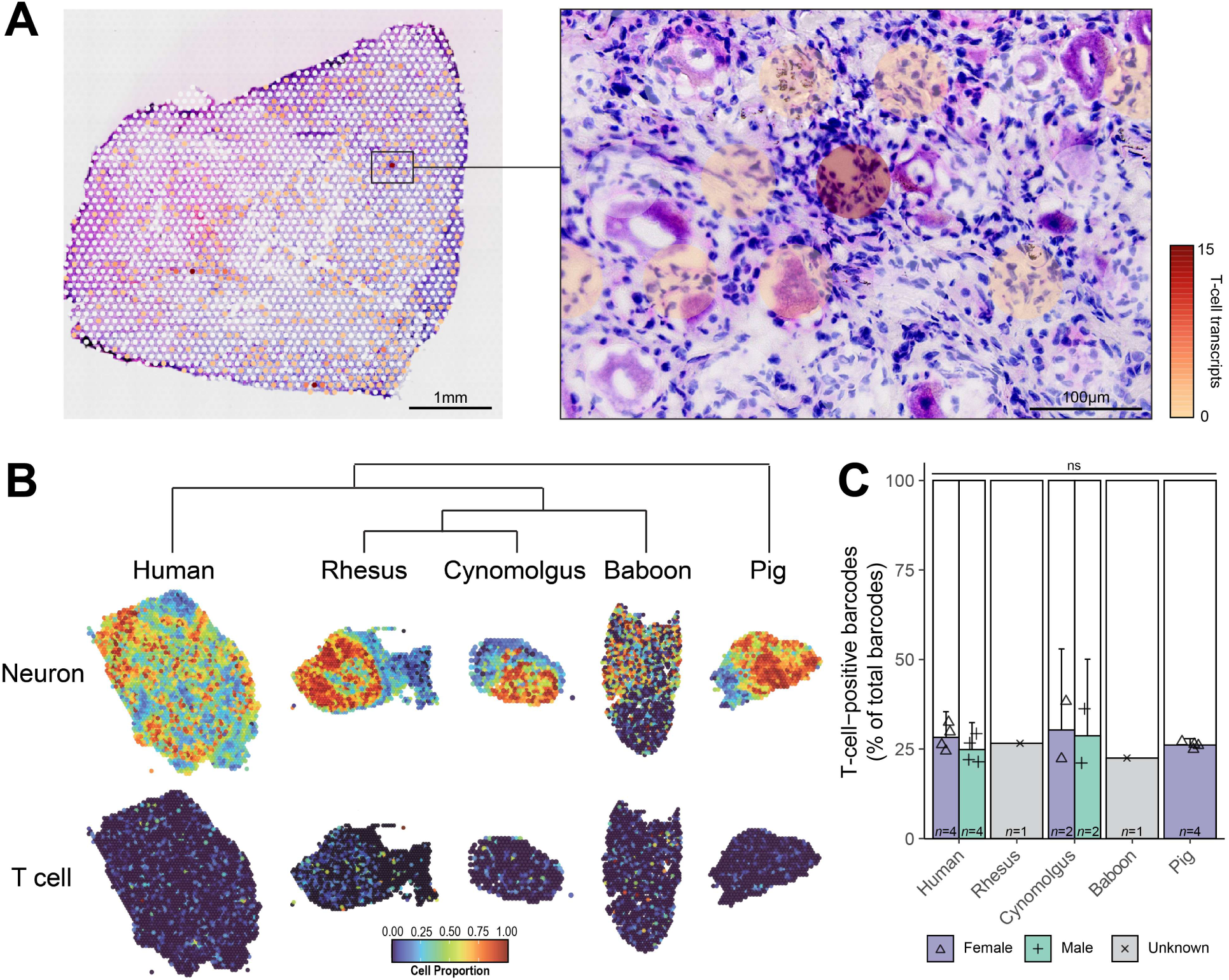
Cell type deconvolution of dorsal root ganglia spatial RNA-sequencing shows similar predicted T-cell signature frequencies across humans and non-human large mammals. **(A)** Frequency of a manually curated list of 10 T-cell marker genes – including CD3 genes and T-cells-specific transcription factor genes – per barcode overlaid over hematoxylin and eosin staining of the human DRG. T-cell genes were spread across the DRG with select high-expressing barcodes overlapping with regions of high nuclear density. **(B)** Representative heatmaps of neuron and T-cell density between species as determined by algorithmic cell type deconvolution. T-cells are predicted to be interspersed throughout the DRG of all species investigated. The dendrograph represents the evolutionary relationship between the included species. Tissue sizes are not to scale. **(C)** Quantification of the proportion of T-cell-positive barcodes shows no differences in predicted T-cell density between species (two-way ANOVA, species × sex *F_1,11_* = 0.07, *P* = 0.80, species main effect *F_2,11_* = 0.71, *P* = 0.51, sex main effect *F_1,11_* = 0.79, *P* = 0.39). ns: not significant. Numbers below columns denote group *n*.

These preliminary results justified more systematic analysis using an unbiased and unsupervised approach to enable robust comparison with other species. To achieve this, we deconvoluted the spatial sequencing data from human, non-human primate, and pig using a single-nucleus RNA sequencing atlas of the DRG to estimate the relative abundance of T-cells and other cell types within each 55 µm spatial barcode. The predicted T-cell proportion within each barcode generated a bimodal distribution, allowing for the designation of barcodes as T-cell-positive or T-cell-negative (**Supplemental Figure 1**). This confirmed that T-cells are present in all DRG niches, including in the neuron-rich bulb and the myelinating Schwann cell-rich nerve root, in all species investigated (**Figure 3B**). The percentage of T-cell-positive barcodes was similar between species, with approximately 25% of barcodes containing a T-cell gene signature. Comparison of human female to male T-cell proportions revealed no significant difference, though the proportion in female DRGs was more variable and trended marginally higher (**Figure 3C**). T-cells are therefore robustly present in the DRG of several evolutionarily divergent large mammals living under different environmental conditions.

### T-cells are abundant in the DRG of large mammals but not rats or mice

To validate the sequencing findings with a more direct cellular quantification method and to enable comparison with smaller mammals such as rodents, we performed immunofluorescence staining on tissue sections from human, rhesus macaque, cynomolgus, pig, rat, and mouse DRG to visualize and quantify CD3^+^ T-cells. This analysis showed that T-cells are interspersed within the DRG parenchyma of human, rhesus macaque, cynomolgus, and pig, and in each of these species were frequently found adjacent to peripherin (PRPH)^+^ neuronal cell bodies (**Figure 4A**). In these large mammals, T-cells were typically excluded from the satellite glia barrier, but some were observed to breach the barrier, which has been previous reported in mouse models of neuropathy [81]. Quantification of CD3^+^ cells confirmed that T-cells were present at similar densities in adult humans and juvenile pigs. No significant sex differences were noted, but the variability was appreciably higher in females, with half of female samples having a higher density than the maximum observed in males. In rats, T-cells were less abundant than in large mammals but were present at rates of approximately 1-2 cells per tissue section on average. In contrast, T-cells were very rare in mice, with only one T-cell detected across all analyzed sections (**Figure 4B**). These results show stark baseline differences in DRG T-cell abundance between large mammals and laboratory rodents, particularly mice.

**Figure 4.**
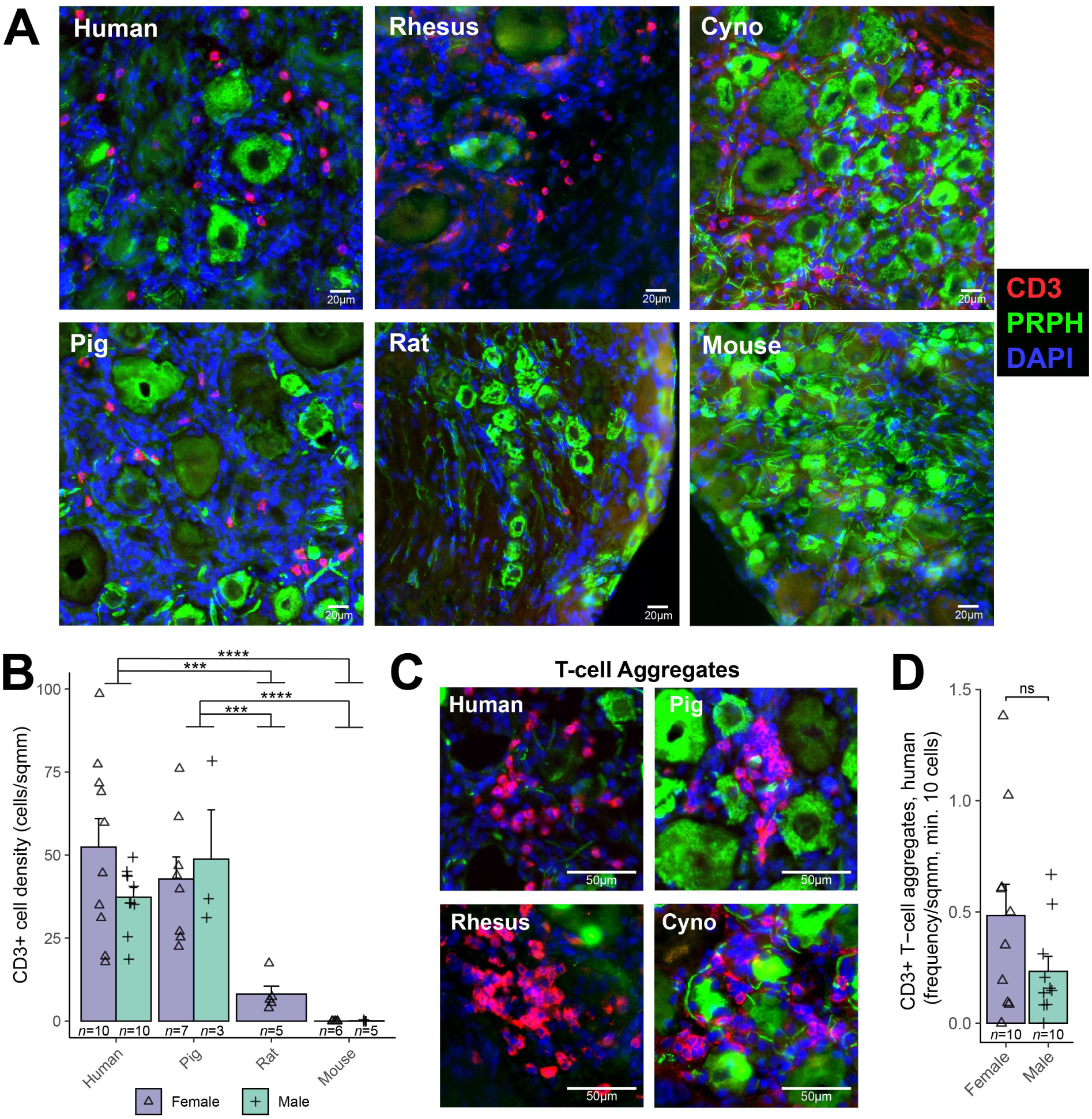
T-cells are robustly present in DRG at similar levels in humans, non-human primates and pigs. T-cells were less frequent in naïve rats and largely excluded from the DRG in mice. **(A)** Representative immunofluorescence photomicrographs of DRG parenchyma T-cells (CD3, red) at 20X magnification. T-cells were present in human, pig, rhesus macaque, cynomolgus, and rat, but were rare or absent in mouse. Neuronal cell body and axonal staining with peripherin (PRPH, green) showed that many T-cells are in close proximity to neuronal cell bodies. DAPI nuclei counterstain in blue. **(B)** Quantification of CD3^+^ T-cell density, expressed in cells/mm^2^, showed a significant effect of species (two-way ANOVA, species × sex; species main effect *F_2,35_* = 18.5, *P* < 0.0001). Rat DRG T-cells were significantly less frequent than in human DRG (*post hoc* pairwise *t*-test, *P_adj_* = 0.0004) or pig DRG (*P_adj_* = 0.0008). Mouse DRG T-cells were also less frequent than in human or pig DRG (both *P_adj_* < 0.0001) or pig DRG (*P_adj_* = 0.0008) but was not significantly different to rat (*P_adj_* = 0.77). The effect of sex was not statistically significant (sex main effect *F_1,35_* = 1.40, *P* = 0.25), though five out of ten adult human female DRGs had higher T-cell density than the maximum in males, with one female doubling the highest male density. Mean ± standard deviation: human female 52.6 ± 27.0; human male 37.2 ± 9.30. Numbers below columns indicate group *n*. **(C)** Representative immunofluorescence photomicrographs of T-cell aggregations observed in large mammal DRGs adjacent to sensory neurons and occasionally breaching the satellite glial barrier. **(D)** In humans, T-cell aggregations were not significantly different between females and males (Welch’s unequal variances *t*-test, *t_12.94_* = 1.61, *P* = 0.13), though again the distribution was wider in females. Numbers below columns denote group *n*. ns: not significant; \*\*\*\**P* < 0.0001.

DRG T-cells occasionally organize themselves into tight clusters or aggregations. In all large mammals, these aggregations of up to dozens of T-cells in the 20 µm section were occasionally observed within the DRG parenchyma; these were typically adjacent to neurons (**Figure 4C**). In humans, these aggregations were not differently abundant between females and males, but again this was more variable and trended higher in females (**Figure 4D**).

### T-cells are present in human DRG early in life

We next investigated whether T-cell density or distribution was altered throughout the human lifespan. First, to explore the presence of DRG T-cells in early life, we performed spatial transcriptomics and cell type deconvolution analysis on lumbar DRGs from a 2-month-old human female and a 14-month-old human male. This showed the presence of T-cell-positive barcodes throughout the DRG, though unlike adults T-cells were more frequent in the white matter and connective tissue within the bulb and less frequently adjacent to neurons (**Figure 5A**). Whole-section quantification found the robust presence of T-cell-positive barcodes in the infant DRGs, but there were fewer than in the adult DRGs (**Figure 5B**).

**Figure 5.**
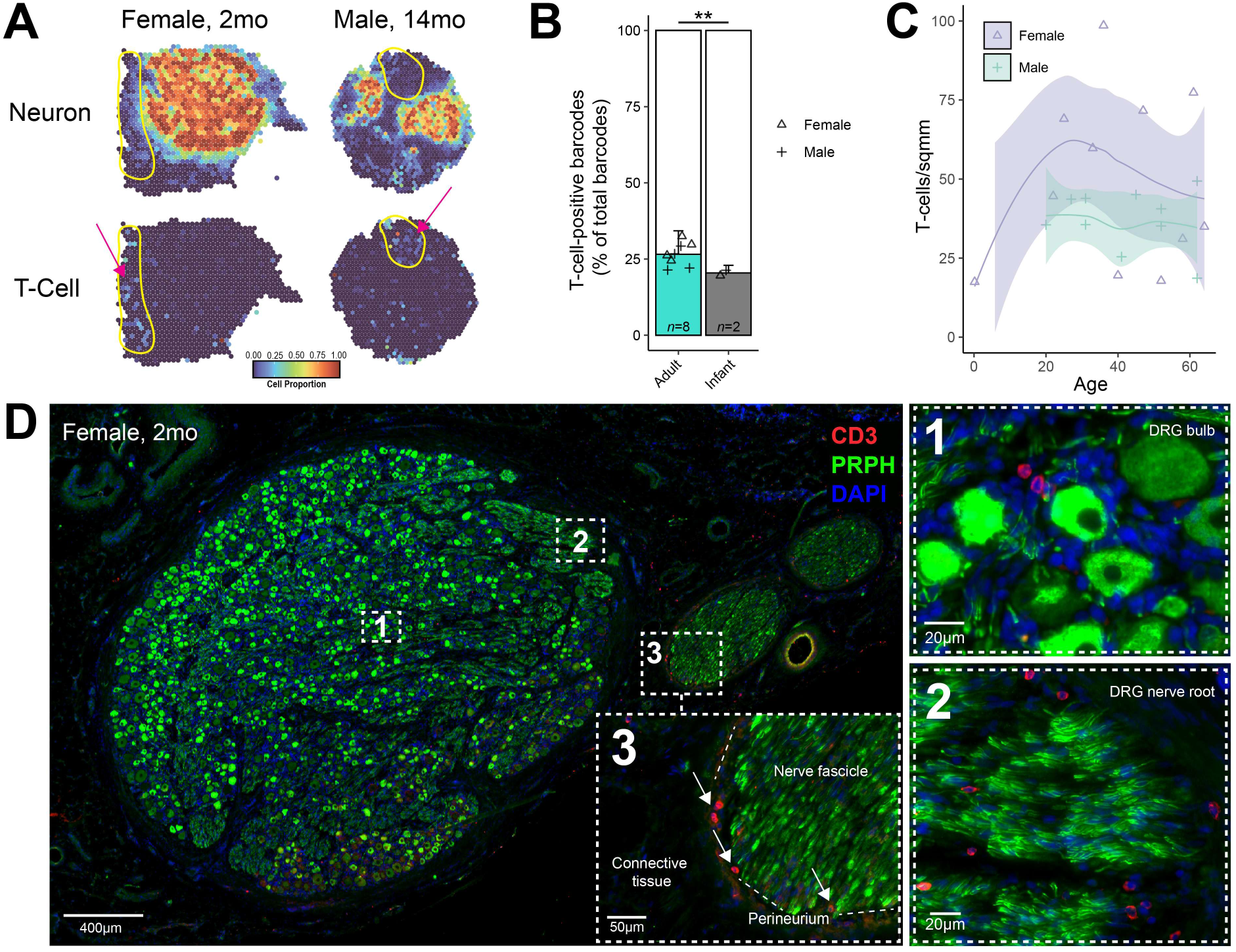
T-cells are present in human infant DRGs, but at lower levels than adults, and may vary in density across the female lifespan. **(A)** Cell type deconvolution of neuron and T-cell spatial transcriptomic signatures in two human infant DRGs. T-cell signatures were robustly detected within the bulb but were strongest in the superficial connective tissue external to the DRG endoneurial space (indicated by magenta arrows). **(B)** There were fewer T-cell-positive barcodes in the infant DRG compared to the adult DRGs in the spatial transcriptomic data (*t_6.28_* = 3.75, *P* = 0.009). Numbers below columns indicate group *n*. **(C)** T-cell density in the human DRG across the female and male lifespan based on immunofluorescence cell counts. Male DRGs had a consistent T-cell density through adulthood. Female DRGs were more variable in the frequency of T-cells; they appeared to trend highest in early adulthood and then decrease with age. The DRG T-cell frequency of the female infant donor is included for comparison. The shaded area represents the 80% confidence interval. **(D)** Representative photomicrographs of a 2-month-old female DRG section staining for T-cells (CD3, red) and sensory neurons (PRPH/peripherin, green). Nuclei are stained with DAPI (blue). Insert (1) shows T-cells near neurons in the DRG bulb. Insert (2) shows T-cells in the DRG nerve root in association with peripherin-immunoreactive axons. Insert (3) shows T-cells present in the perineurium of a nerve fascicle adjacent to the DRG bulb. T-cells were otherwise absent from these fascicles in the analyzed sections. \*\**P* < 0.01.

T-cell frequency was then evaluated with respect to age across all human DRGs. This confirmed the presence of T-cells in infant DRG at rates lower than in adult DRG. Furthermore, a possible sex-dependent effect emerged whereby T-cell density was consistent across adulthood in males, but in females was highest in early adulthood and decreased toward male levels in late adulthood; however, despite this trend, there was still substantially more variability in T-cell frequency in females than in males that was not solely explained by age (**Figure 5C**).

Immunofluorescence staining in the 2-month-old female DRG showed CD3^+^ T-cells in proximity to neurons, but these were sparser than in the adult samples. However, a similar density to adults was found in the white matter within the DRG. Nerve fascicles external to the DRG contained very few T-cells in the sections investigated, but they were frequently noted within or adjacent to the fascicle perineurium and in both endoneurial and external connective tissue, which is consistent with the transcriptomic findings (**Figure 5D**). Overall, while T-cells remain numerous in human infant DRG, they are less frequent than in adults, appear less likely to associate with neurons, and were preferentially distributed elsewhere, including in the DRG white matter, perineurium, endoneurium, and internal and external connective tissue.

### Altered spatial and intercellular associations of human DRG T-cells across sex and age

The stark qualitative differences observed in T-cell distribution in the human infant DRG prompted us to quantify the spatial associations between T-cells and other DRG cells in more detail. To do this, we leveraged the more comprehensive cell-type coverage offered by the spatial transcriptomics data by performing an exploratory analysis of T-cell colocalization with the 13 other DRG cell types identified by cell type deconvolution and performed comparisons within the human adult and infant DRG and between females and males. Principal components analysis was performed on Pearson correlations between T-cells and other cell signatures. One principal component separated infant from adult DRGs (PC1, 30.4% of variance) with the highest loadings contributed by endothelial cells (0.42), smooth muscle cells (0.36) and macrophages (0.35). Females showed more variability in the PCA than males. We hypothesized that there may be a sex hormonal influence on T-cell retention, so we attempted to account for this variation by separating females into post-teen pre-menopausal (34-44 years) and post-menopausal (53-57 years) adult females. PC2 (22.9% of variance) provided separation between pre-menopausal and post-menopausal females, the latter grouping more robustly with males, with the highest loadings originating from granulocytes (0.55), fibroblasts (0.48), and myelinating Schwann cells (0.32; **Figure 6A**).

**Figure 6.**
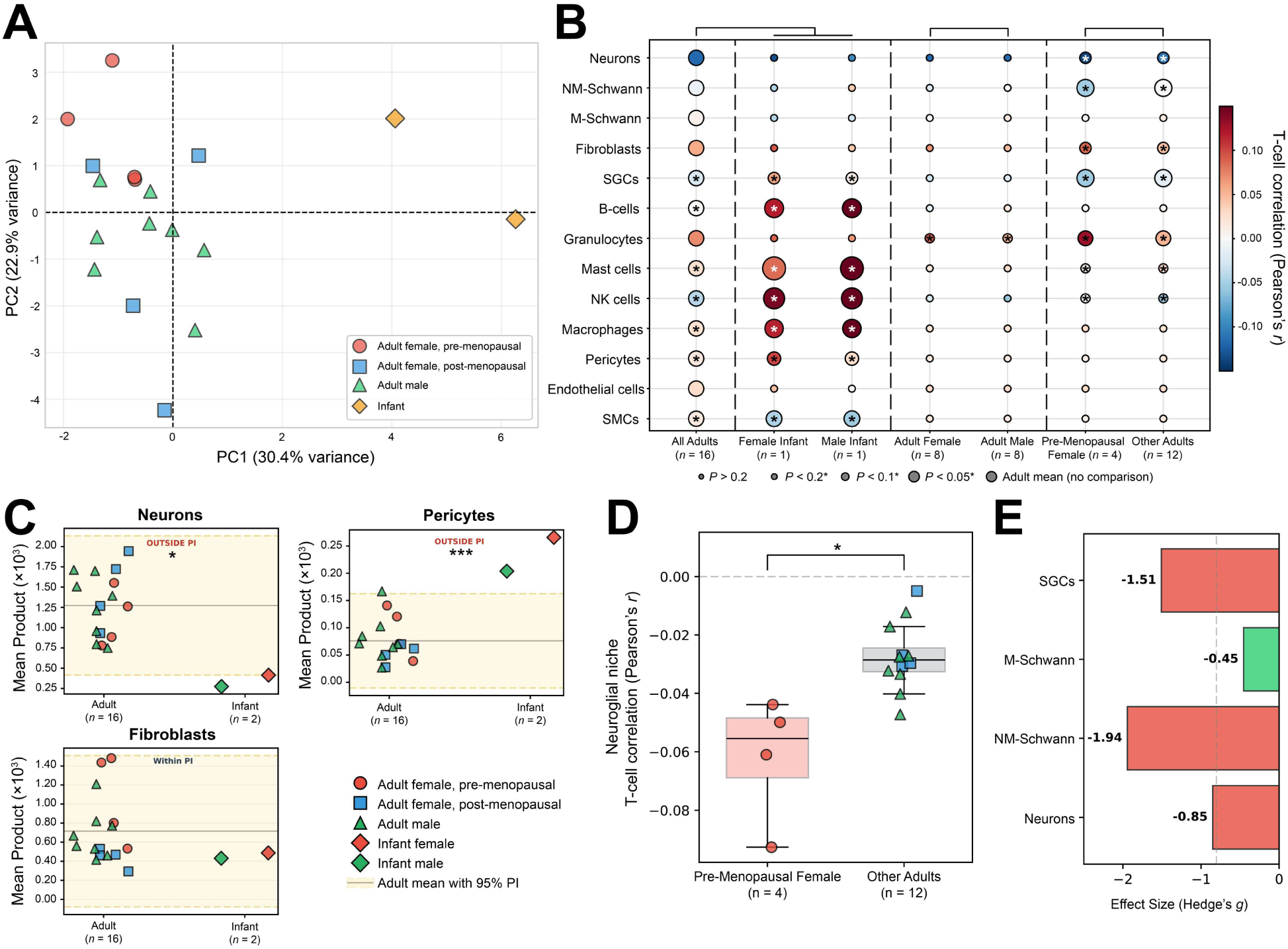
Human DRG T-cells display altered intercellular relationships across sex and age. **(A)** Principal components analysis plot of T-cell colocalization scores with other DRG cell subpopulations. **(B)** Dot plot summarizing sample-level Pearson’s *r* correlation scores between T-cells and 13 other human DRG cell types. Dots are colored by Pearson’s *r* and sized by p-value. Text within or adjacent to dots denotes the Pearson’s *r* value for the given cell type with T-cells; boldened text denotes *P* < 0.2. Dendrograms denote the groups included in each comparison. **(C)** Comparison of mean product colocalization scores between adults and infants showed outside the 95% prediction interval. T-cells colocalized less with neurons (*z* = -2.38, *P* = 0.036), more with mural cells including pericytes (*z* = +4.04, *P* = 0.001), and were no different in their spatial relationship with fibroblasts (*z* = - 0.71, *P* = 0.50). **(D)** Pearson correlation between T-cells and the neuroglial niche (neurons, satellite glial cells, and myelinating and non-myelinating Schwann cells) showed a significantly reduced association between this niche and T-cells in post-teen pre-menopausal females compared with other adults (Bonferroni-corrected Mann-Whitney *U* test, Hedges’s *g* = -2.27, *P_adj_* = 0.042). **(E)** The contributing cell types to the decreased T-cell-neuroglial association were non-myelinating Schwann cells, satellite glial cells, and neurons, but not myelinating Schwann cells as determined by a large effect size threshold of Hedges’s *g* ± 0.8 (boldened text; red bars denote exceeding effect size threshold). M-Schwann: myelinating Schwann cells; NM-Schwann: non-myelinating Schwann cells; PI: prediction interval; SGC: satellite glial cell; SMC: smooth muscle cell. \**P* < 0.05, \*\*\**P* < 0.001.

Detailed analysis of these Pearson correlations showed several changes between adult and infant DRGs, and though there were few differences between female and male DRG overall, differences emerged when comparing post-teen pre-menopausal females to post-menopausal females and adult males (**Figure 6B; Supplemental Table 2**). To assist with assessment of meaningful differences between the small infant sample and adults, the infant DRG mean product colocalization scores were compared against the 95% prediction interval (PI) of adult DRGs showed that in infants T-cells had decreased colocalization with neurons, increased colocalization with pericytes and mast cells, but fell well within the adult distribution for fibroblasts and other cell types (**Figure 6C**; complete results in **Supplemental Figure 2** and **Supplemental Table 3**). Finally, to further link spatial alterations in cell type changes to distinct tissue neighborhoods, composite colocalization scores were created for cells contributing to neuroglial, stromal, inflammatory, and vascular niches. T-cells showed significantly reduced colocalization with the neuroglial niche in pre-menopausal females compared to other adult DRGs (**Figure 6D**). The strongest individual contributors to this association were non-myelinating Schwann cells, satellite glial cells, and neurons, but not myelinating Schwann cells (**Figure 6E**). Given that myelinating Schwann cells are more commonly distributed toward nerve fiber rather than neuronal cell body-rich DRG regions, this together suggests reduced T-cell-neuron associations in the pre-menopausal female DRGs investigated.

We validated this reduced neuroglial colocalization in pre-menopausal female DRG by training a random forest classifier to discriminate pre-menopausal females from all other human adults based on the T-cell colocalization patterns of the 13 other cell types. This produced a leave-one-out cross-validation accuracy of 87.5% with a specificity of 91.7% (11/12 true negatives correctly identified) and a sensitivity of 75% (3/4 true positives correctly identified). This further supports a robust multivariate signature of T-cell spatial localization that is specific to pre-menopausal female adults.

### Human DRG T-cells express tissue residence markers

We then sought to phenotype these DRG T-cells, primarily to determine marker expression patterns consistent with tissue retention and Trm status. Flow cytometry analysis of enzymatically dissociated human DRGs from a 29-year-old male showed that the majority of DRG T-cells are CD8^+^, outnumbering CD4^+^ T-cells approximately 3-to-1, though some interindividual variability should be anticipated (**Figure 7A**). The activation and tissue retention marker CD69 was expressed by 85.7% of CD4^+^ T-cells and 94.8% of CD8^+^ T-cells. CD103, which is typically associated with Trms localized to barrier sites, was highly expressed in approximately 40% of CD8^+^ T-cells but only at low expression levels in CD4^+^ T-cells (**Figure 7B-C**). Immunofluorescence analysis of tissue sections recapitulated a Trm phenotype (**Figure 7D**). As an isoform of CD45, CD45RO expression confirms that these CD3^+^ T-cells are hematopoietic in origin and further distinguishes them from CD3^lo^ perineuronal cells. Taken together, the majority of DRG T-cells express a phenotype consistent with Trm status, with the specific integrins promoting local retention potentially differing between CD8^+^ and CD4^+^ cells.

**Figure 7.**
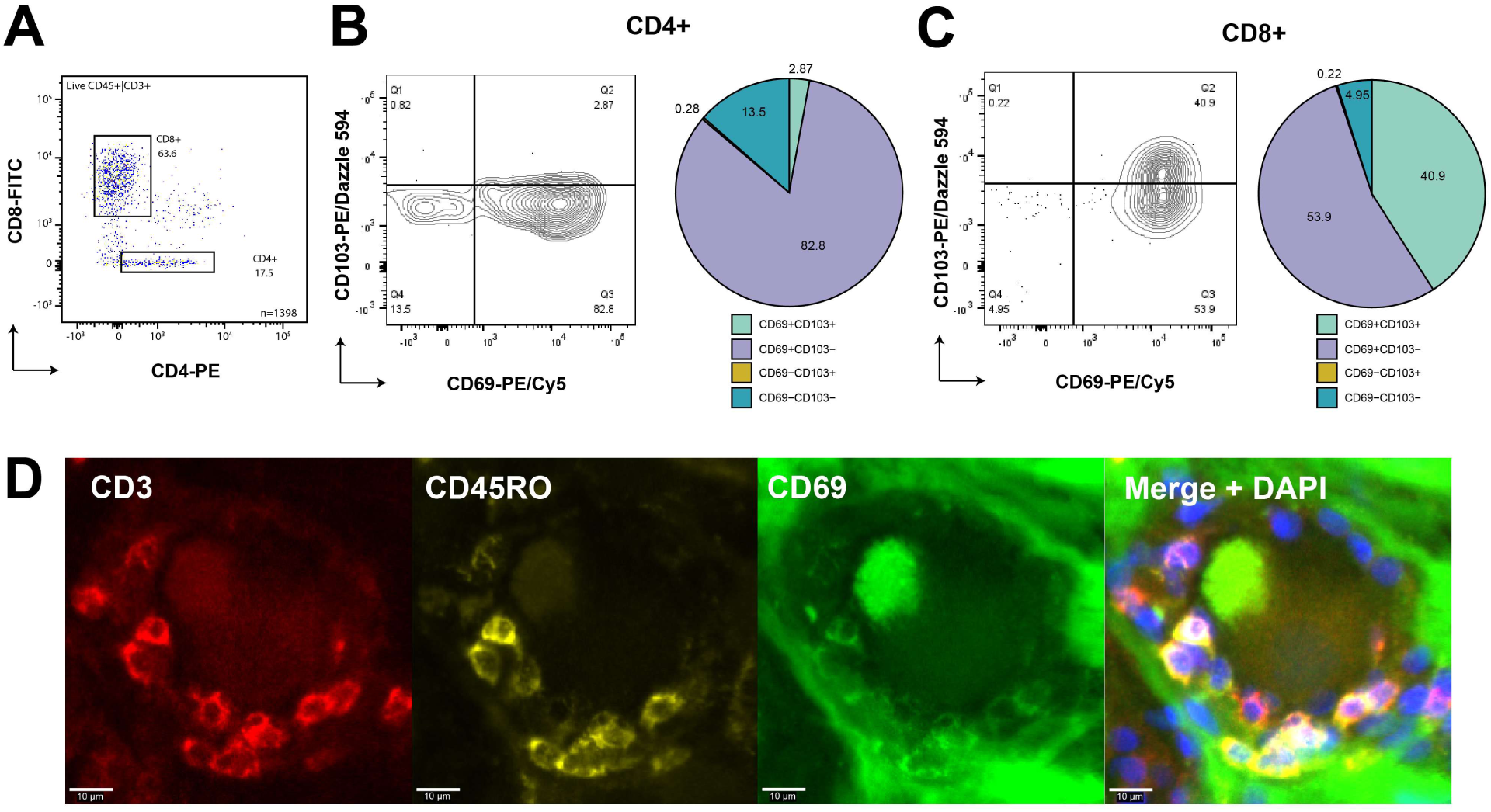
Human DRG T-cells express a Trm phenotype. **(A)** CD8^+^ and CD4^+^ DRG T-cells from a 29-year-old donor. CD8^+^ T cells constituted the majority of CD45^+^CD3^+^ cells. **(B)** Biaxial plots and quantification showing altered residence marker expression of CD69 and CD103 in these CD4^+^ and **(C)** CD8^+^ DRG T-cells. **(D)** Representative DRG immunofluorescence photomicrographs of key Trm markers CD45RO and CD69 *in situ,* focusing on a cluster of T-cells breaching the satellite envelope of a sensory neuron. The expression of CD45RO and CD69 on a majority of DRG T-cells is consistent with a Trm phenotype. 20X images are digitally zoomed for 60X final magnification.

## Discussion

Here, we have shown that the presence of DRG T-cells within molecular features associated with tissue residency is conserved among humans, non-human primates, and pigs without a known history of peripheral nervous system disease. In contrast, T-cells were infrequent in naïve rat DRG and largely absent from naïve mouse DRG. In humans, no clear sex difference in T-cell density was delineated, though we provide evidence of higher variability in T-cell number in females which may be elevated in pre-menopausal adulthood, and they alter their spatial distribution and interactions with other cells in an age- and sex-dependent manner. We have also shown that T-cells are present in the dorsal root ganglion of large mammals in the early weeks of life. In human infancy, these T-cells were occasionally observed near neuronal cell bodies but preferentially occupied DRG epi-and perineurial spaces. Finally, we demonstrated that these human DRG T-cells express canonical residence markers, confirming a Trm phenotype consistent with investigations in human trigeminal ganglion [75].

### Species diferences between mice, rats, and large mammals

We observed stark differences between T-cell numbers in the DRG of large animals compared to rodents. Laboratory rodents continue to be workhorse models of *in vivo* human biological processes and have led to tremendously important insights into immunology and neuroscience [45; 49]. However, differences between rodents and humans have drawn attention to potential limitations of these models, such as the absence in mice of microglial-like perineuronal macrophages that are found in human and other large mammal DRGs [82]. Understanding whether the observed difference in T-cell residency is an inherent species difference or the by-product of environmental parameters is of great importance to clarifying how to interpret the findings of many papers studying the peripheral nervous system and associated disease.

Our findings provide evidence against some explanations of this species difference. Firstly, the robust presence of T-cells in DRG is not a human- or primate-specific phenomenon and instead appears to be evolutionarily conserved across the large mammals we examined. Second, humans and pigs were assessed with comparable chronological age to the rodents, which suggests that developmental influences cannot fully explain the observed difference.

One feasible environmental cause for the observed paucity of DRG T-cells in rodents is that they were raised and housed in specific pathogen-free (SPF) conditions. This has long been proposed as a potential confounder of rodent experiments because the lack of exposure to usual levels of pathogenic and innocuous antigens in laboratory rodents contributes to immunological naivety [16; 45]. The immune systems of laboratory mice, as measured by splenocyte phenotyping, are less mature than wild mice, particularly with respect to T-cells [1]. While a heritable contribution of selection pressures following generations of selective breeding cannot be ruled out, evidence that rewilding laboratory mice induces maturation of adaptive immunity and individualized clonal diversity suggests a stronger nonheritable influence [11; 17; 54]. Importantly, the large non-human animals included in this study were also laboratory animals housed under controlled conditions, which may suggest that these differences are not fully explained by housing conditions.

The findings reproduce previous reports of an infrequent but consistent T-cell presence within naïve rat DRG. In two papers from the same group, very few (1-2 cells per section) TCR^+^ T-cells were observed in naïve rat DRG; interestingly, the authors note that the T-cell density in naïve non-SPF (conventionally-housed) male Sprague-Dawley rats is approximately triple that of naïve female Wistar rats maintained in SPF conditions [27; 28]. Though strain and sex may play a role here, antigen exposure history emerges as a likely contributor. Additional studies in rats similarly report low levels in naïve states [46; 63]. In naïve mice, T-cells are robustly present in a single-cell RNA sequencing data set of sympathetic ganglia, but in somatosensory ganglia the presence of T-cells differs between investigations [9; 44]. This may reflect differences in tissue preparation protocols since it is possible that lymphocytes within the DRG meninges and other extraneous tissues may contribute to detections within such studies. Evidence exists to suggest that T-cells may infiltrate the DRG of aged mice, but appear to express an infiltrating rather than primary residence phenotype [86]. In one study, immunofluorescence and flow cytometry were used to detect T-cells in naïve 8-week-old C57BL/6J mouse DRG, finding increased numbers of regulatory T-cells in female compared to male and ovariectomized female DRGs indicative of a sex hormonal contribution [21]. To clarify these conflicting reports, *in situ* investigations comparing laboratory rodents with true wild or ‘dirty’ rewilded rodents will be valuable [3].

### DRG T-cell spatial distribution and residence patterns

DRG T-cells were observed to form large aggregates, sometimes comprising dozens of cells, within the endoneurial space in most large mammal samples investigated. Similar aggregations have been noted in rats following nerve injury, but not in naïve states [28]. In some cases, T-cells appeared to breach the perineuronal satellite glia barrier in a manner similar to what has been reported in human trigeminal ganglion [78] and in a murine model of chemotherapy-induced neuropathic pain [81]. The functional relevance of these aggregates in health and disease is worthy of further investigation given their frequency.

Human DRG T-cells also largely express residence markers. CD103 (integrin αE), which was predominantly identified on CD8^+^ T-cells, binds E-cadherin [39], which in human DRG is enriched in *MRGPRX1*-expressing C-fibers associated with itch [8], and in satellite glial cells [9]. This not only confirms a Trm phenotype in most T-cells – consistent with prolonged local retention and repeated opportunities to influence the DRG milieu – but also raises the possibility of an integrin-mediated mechanism for localized T-cell organization within DRG to direct responses toward specific neurons or tissue microenvironments. Given that the Trm compartment is known to be dynamic in other tissues and is strongly shaped by environmental exposures such as injury and infection, this possibility requires further investigation.

### DRG T-cells show altered spatial distribution in pre-menopausal women

Sexual dimorphism in interactions between T-cells and sensory neurons has been proposed as an influential modulator of pain processing [51; 52; 68]. One possibility for our observed differences in T-cell spatial distribution in females is variation in sex hormones across the lifespan, which affect T-cell phenotype in animal models [47] and in transgender individuals undergoing hormone replacement therapy [36; 59]. Some female donors included in our results had a history of hysterectomy, birth control use, and other factors that may modulate sex hormones, but we did not observe any remarkable influence of these factors on DRG T-cell frequency in this sample. Further investigation appears warranted, particularly given the overrepresentation of pain and peripheral nervous system disease in older women [37].

### DRG T-cells and age

Infancy is a key period for immune education, particularly for the generation of Trm as they are seeded and retained in sites of previous infection [53; 84]. These observations suggest that DRG T-cell seeding has already commenced, but is yet incomplete, as early as the second postnatal month. Later in life, adaptive immune cells typically become functionally impaired with the onset of senescence or ‘inflammaging’ [50; 67]. We did not find evidence of increased T-cell colonization of DRG in older individuals, but this does not rule out functional alterations that may drive age-associated inflammation.

### Limitations

This study has some limitations. The sample size is appropriate for species-level comparisons, but the high interindividual variability when measuring along the dimensions of sex and age in human adults should render these analyses exploratory. While all individuals included had no recorded history of peripheral nervous system disease or injury, we cannot account for a lifetime of infectious exposures that may impact DRG T-cell retention and contribute to this variability. We can only speculate on the functional relevance of the reported T-cell aggregations and T-cell-sensory neuron spatial interactions. Finally, while DRG T-cells express canonical Trm markers, we cannot conclusively determine long-term non-recirculating residency. We believe the results reported here provide strong justification for additional investigation into these observations.

## Conclusions

The presence of T-cells in the DRG of large mammals throughout the lifespan raises important questions about how they may engage in bidirectional interactions with sensory neurons both in maintaining health and contributing to peripheral nervous system pathologies including viral infection, neuropathy, and chronic pain. T-cell interactions with sensory neurons are important to many biological processes across organ systems [12; 26; 56; 72], therefore clear reporting of housing conditions and potential opportunities for infection exposure are key to the interpretability of rodent models investigating these interactions where immune naivety may confound results. To address these concerns, large mammal and wilded rodent models [22] may emerge as particularly informative complementary models of somatosensory nervous system disease and injury as they may more closely recapitulate the human reality. These models will be crucial to better understanding the functional relevance of T-cells to peripheral nervous system health and disease as failure to account for baseline immune residence in the DRG may lead to systematic misinterpretation of neuroimmune mechanisms in rodent disease models.

## Acknowledgements

We are grateful to the organ donors and their families whose gift made possible the human aspect of this work, to the Southwest Transplant Alliance whose partnership enabled organ donor tissue collection, and to all members of the UT Dallas human sensory ganglion collection teams. We thank Merck, particularly Darrell Henze and Becky Klein, for access to rhesus macaque tissue, Ulrike Hoffmann (UT Southwestern, TX) for pig tissue, Muhammad Saad Yousuf and Brodie Woodall (UT Dallas, TX) for rat tissue, and the staff of Deep Cuts (Richardson, TX) for access to pig lymph node tissue. We also thank Derek Howard, Shreejoy Tripathy, and Angelika Lampert for assistance with pig gene homology questions. The microscopy and flow cytometry data was made possible through the equipment and technical expertise of Joe Lombardo of the UT Dallas Microscopy Core Research Facility and Jacob Henderson of the UT Dallas Flow Cytometry Core Research Facility respectively. Figure 1 was created in BioRender: https://BioRender.com/ldhzub4.

This work was funded by the National Institutes of Health grant U19NS130608 to TJP.

## Declarations

### Conflict of interest statement

TJP is co-founder of 4E Therapeutics, PARMedics, NuvoNuro, Nerveli, and Ted and Greg’s.

### Data availability statement

Non-human primate dorsal root ganglion spatial transcriptomic raw data is publicly accessible on GEO (accession number GSE313530). Infant human dorsal root ganglion spatial transcriptomics raw data is accessible via SPARC. Other analyzed spatial transcriptomics data sets have been previously published elsewhere and are cited in the manuscript. Original code is available on the Open Science Framework (DOI: 10.17605/OSF.IO/2FB9Y). Raw microscopy files will be made available upon request to the corresponding author. Other data are available in the supplemental materials.

## Methods

### Sample Information

Human DRGs (*n* = 64 from 52 unique donors) were collected post-mortem from organ donors through the Southwest Transplant Alliance (Dallas, TX, USA). Procedures for human tissue procurement were approved by the University of Texas at Dallas Institutional Review Board (MR-15-237). Informed consent from the donor and/or their legal next of kin was obtained by the Southwest Transplant Alliance. Information on donor age, sex, ethnicity, cause of death, positive serologies, and medical history as part of the donor risk assessment inventory and available medical records were collected as part of standard organ procurement procedures. DRGs were extracted and either rapidly frozen in crushed dry ice within 4 hours of cross-clamp and stored at -80°C long-term in line with previously published protocols [66] (spatial transcriptomics, immunofluorescence), or maintained either in artificial cerebrospinal fluid or supplemented Hibernate-A medium as previously described [38] at 4°C until tissue dissociation (flow cytometry). Human lymph node was similarly extracted and frozen for use as an immunostaining positive control.

Sixteen fresh-frozen lumbar cynomolgus monkey DRGs from four individuals (*Macaca fascicularis*; two males and two females) aged 5–10 years were collected from the lumbar region. Animal housing, euthanasia, and tissue collection were performed by Worldwide Primates, Inc. Tissues were shipped to the University of Texas at Dallas on dry ice and stored at −80°C until processing.

One fresh-frozen dorsal root ganglion each from one adult rhesus macaque (*Macaca mulatta*, lumbar level) and one adult olive baboon (*Papio anubis*, cervical level) were collected at necropsy following humane euthanasia under deep anesthesia. Tissues were rapidly harvested, trimmed, and immediately frozen on aluminum foil lining a metal plate on dry ice. Frozen tissues were then shipped to UT Dallas overnight on dry ice and stored at −80°C until processing.

Pig (*Sus scrofa,* German Landrace, wild type, no backcrossing) spatial transcriptomic data was reanalyzed from a previously published data set [33]. Pig DRG tissue used for immunofluorescence staining was provided by Dr. Ulrike Hoffman at the University of Texas Southwestern Medical Center (Dallas, TX, USA), which were frozen in crushed dry ice and stored at -80°C until processing, and from archival sections made available by Grünenthal (Aachen, Germany) stored at −80°C. Pig lymph node tissue used as a positive control for immunostaining was obtained from Deep Cuts (Richardson, TX), where it was taken from 4°C storage on the same day as tissue processing, frozen in crushed dry ice, and thereafter stored at -80°C until processing.

Female Sprague-Dawley rats (*Rattus norvegicus*; Envigo, IN, USA; *n* = 5, aged 21 weeks) and female (*n* = 6) and male (*n* = 5) C57BL/6J mice (*Mus musculus*; Charles River, MA, USA; aged 6 or 12 weeks) were obtained and handled in accordance with approved University of Texas at Dallas Institutional Animal Care and Use Committee protocols 19-08 and 14-04 respectively. Rodents were group-housed under specific pathogen-free conditions in a 12-hour standard light-dark cycle and had access to water and standard chow *ad libitum*.

Rodents were placed under deep isoflurane anesthesia; rats were then euthanized by rapid decapitation and mice by cervical dislocation. DRGs and spleens were extracted, immediately frozen in optimal cutting temperature compound (Sakura), and stored long-term at -80°C. Rats and mice did not undergo any interventional procedures prior to euthanasia.

All animal care and experimental procedures were approved by the institutional IACUC and conducted in accordance with the *Guide for the Care and Use of Laboratory Animals* and applicable regulatory guidelines. Complete sample information is available in **Supplemental Table 4.**

### Flow cytometry

Flow cytometry staining for CD45, CD11b, and CD3 (panel 1) was performed on DRGs from 20 human donors, while staining for T-cell residence markers (panel 2) was performed on DRGs from a single donor. Fresh human DRGs were trimmed, cut into 1 mm pieces, and dissociated in 2 mg/mL Stemxyme 1 (Worthington LS004107; panel 1) or 2 mg/mL collagenase IV (Worthington LS004188; panel 2) and 4 mg/mL DNase I (Worthington LS002139) in Hanks’ balanced salt solution (HBSS; Gibco 14170-112) at 37°C in a shaking water bath. Mechanical trituration was performed at regular intervals with a glass Pasteur pipette. Per-sample dissociation details and results are provided in **Supplemental Table 5**.

The resulting cell suspension was filtered through a 70 µm nylon mesh strainer, spun at 350 *g* for 5 min at room temperature (parameters used for all subsequent spin steps), and the supernatant was aspirated. Cells were incubated in 1 mL red blood cell lysis buffer (Biolegend 420301) for 10 min at room temperature in the dark, then underwent myelin debris removal using magnetic anti-myelin beads (Miltenyi 130-096-733) in LS columns (Miltenyi 130-042-401) per the manufacturer’s recommendations. Cells then underwent live/dead staining with 2 µL of Zombie-UV viability dye (Biolegend 423107) in 1X phosphate-buffered saline (PBS) for 15 min at room temperature. Cells then underwent Fc blocking with 5 µL of human TruStain FcX blocking solution (Biolegend 422302) in 200 µL of cell staining buffer (Biolegend 420201) for 10 min at room temperature. Primary antibody mix (**Supplemental Table 6**) was then added and incubated for 30 min on ice. Cells were then washed once in cell staining buffer. Single-stained compensation controls were performed by staining 1 drop of beads (ThermoFisher 01-3333-42) with 1 µL of primary antibody for 20 min followed by wash and resuspension in staining buffer. Cell sorting was performed on a five-laser BD FACSAria Fusion. The resulting data file was gated and analyzed in FlowJo v10.9.

### Transmission electron microscopy

Human DRG from a single donor (male aged 41 years, spinal level L4) was drop-fixed in 4% paraformaldehyde (PFA) for 24 hours immediately following tissue collection, then washed in PBS. Small pieces of about 1 mm^3^ were postfixed in 10% formalin for 48 hours, washed in PBS and fixed in 2.5% glutaraldehyde in PBS, pH 7.4 at 4°C for 24 hours. The tissue was washed in PBS and transferred into 2% aqueous osmium tetroxide solution for 1 hour. The samples were then dehydrated in an ascending series of ethanol and embedded in epon araldite embedding medium (TAAB Laboratories) at 60°C for 48 hours. Semithin sections with a thickness of 500 nm were prepared and stained with 0.05% toluidine blue in borate buffer. Ultrathin sections of 70-90 nm thickness were cut using an ultramicrotome (Leica), stained with 4% uranyl acetate and Reynolds lead citrate, and examined using a transmission electron microscope (FEI Tecnai 120kV Spirit). An AMT Camera with AMT_V7.0.1 software was used for image capturing. Three sections from three different areas were investigated.

### Spatial transcriptomics

Previously published adult human [74] and pig [33] 10x Visium V1 spatial transcriptomics data were reanalyzed for this study. Non-human primate (10x Visium V1) and human infant (10x Visium V2 CytAssist) data sets are reported here for the first time.

### Spatial transcriptomics acquisition

Tissue embedding, sectioning, and fixation were performed following previously described methods [33] and the 10x Genomics Visium Spatial Gene Expression Demonstrated Protocol (CG000160). Tissue permeabilization on non-human primate was carried out for 12 minutes according to the Visium Spatial Gene Expression User Guide (CG000238 Rev E). Tissue fixation and pre-Visium staining on human infant DRG was performed according to 10x Genomics Visium CytAssist Spatial Gene Expression for Fresh Frozen Demonstrated Protocol (CG000164 Rev A). Human infant Visium data was collected using optimized conditions for Visium CytAssist according to 10x Genomics User Guide (CG000495 Rev E).

Spatial gene expression libraries were generated using the Visium Spatial Gene Expression Reagent Kits (16 reactions; PN-1000186), Library Construction Kits (16 reactions; PN-1000190), and Visium Spatial Gene Expression Slide Kits (16 reactions; PN-1000185). Library preparation followed the manufacturer’s instructions (User Guide CG000239 Rev F). Probe-libraries for the infant human DRGs were generated according to the 10x Genomics User Guide (CG000495 Rev E) using the Visium Human Transcriptome Probe Kit (PN-1000466). Sequencing was performed at the Genome Center at the University of Texas at Dallas using the Illumina NextSeq2000 platform.

Non-human primate Illumina BCL files were processed using the Space Ranger pipeline (v1.1, 10x Genomics) to generate FASTQ files and align sequencing reads to the *Macaca fascicularis* reference genome (Ensembl assemblies: *Macaca_fascicularis_6.0; Mmul_10; Panu_3.0*), while integrating the sequencing data with bright-field microscopy images of the corresponding tissue sections. The raw data has been made available on GEO (accession number: GSE313530).

Sequencing quality control metrics for non-human primate DRGs are reported in **Supplemental Table 7**. Human infant spatial transcriptomics quality control metrics are reported in **Supplemental Table 8**.

### Manual T-cell signature analysis

Initial qualitative analysis estimated the presence of T-cell-related barcodes in Visium barcodes of human DRG by counting those which expressed T-cell-restricted genes within Loupe Browser software (v8.0.0, 10x Genomics). Marker genes for T-cells tend to be conserved between species and include *CD3D/E/G*, T-cell receptor (*TCR*) genes, *ZAP70*, and other modules [15].

### Cell type deconvolution and intercellular analysis

The spatial deconvolution tool SONAR [40] was implemented to predict the cell type proportions in 10x Visium barcodes across 66 DRG sections from human donors and non-human species (**Supplemental Table 4**). As our group has shown previously [42; 65], this technique allows estimation of the contribution of each cell type to the transcriptomes of each spatial barcode. A single-cell RNA-seq dataset of human lumbar DRG [8] was used as a reference to guide deconvolution. The Seurat (v5.0) function FindAllMarkers() [25] was used to identify highly enriched markers (top 250 genes ordered by adjusted *P*-value with log_2_-fold change > 1.0 per cell type) and generate a signature matrix of 2693 genes.

Although some of the human genes did not have orthologues in animal species, there were at least 160 orthologous markers for all 14 cell types per species (**Supplemental Table 9**), well above the 65 markers across 12 cell types for which SONAR has demonstrated effective deconvolution [40]. All spatial barcodes overlapping tissue were processed with the scripts and accessory files provided by SONAR (https://github.com/lzygenomics/SONAR) using R v4.3.3 and MATLAB R2024a. The spot_min_UMI parameter in the SONAR.preprocess() was set to 2 to for all slides to allow processing of the lower sequencing depth *Papio anubis* samples, and default settings were used for all other parameters. The estimated T-cell proportion per barcode generated a bimodal distribution with a cut-off of 10^-4^ determined to separate barcodes with high vs low estimated T-cell proportions. The barcodes with a high predicted proportion of T-cells were designated as T-cell-positive.

#### Colocalization metrics

For each tissue section, T-cell colocalization with the 13 other cell types was quantified using two metrics: (1) Pearson correlation across spatial spots, providing scale-independent spatial co-occurrence, and (2) the mean product of T-cell and target cell-type proportions, to reflect both spatial co-occurrence and marginal abundance of each cell type. Pearson correlation was used to capture the normalized spatial pattern while the mean product is sensitive to colocalization magnitude.

#### Principal component analysis

Principal component analysis (PCA) was performed using the Pearson correlation of cell-type proportions. Separate PCAs were conducted for each metric for the 18 human tissue samples (16 adult + 2 infants), with the first two principal components retained for visualization and interpretation of component loadings.

#### Composite variables

Composite variables were defined based on established cellular compartments and functional roles within DRG, composed of sensory neurons and associated glial cells, immune populations, and stromal/vascular cell types that together form a multicellular microenvironment surrounding neuronal somata [74]. These included neuroglial (neurons, non-myelinating Schwann cells, myelinating Schwann cells, satellite glial cells), immune (B-cells, granulocytes, mast cells, natural killer cells, macrophages), stromal (fibroblasts, pericytes, endothelial cells, smooth muscle cells), and vascular (pericytes, endothelial cells, and smooth muscle cells) composites corresponding to different tissue niches.

#### Statistical analysis

Statistical comparisons were conducted at the sample level (tissue section) and donor level for Pearson correlation and mean colocalization product values. The comparisons were: (1) human adult females vs. adult males (*n* = 8 vs. 8 samples; 4 donors per group, 2 sections per donor); (2) human infants (*n* = 2) vs. adults (*n* = 16); (3) younger adult females (aged < 50 years, range 34-44 years, *n* = 4) vs. all other adults (*n* = 12); (4) Younger adult vs. older adult females (aged ≥ 50 years, range 53-57 years, *n* = 4 vs. 4); and (5) older adult females vs. males (n = 4 vs. 8). Due to the small infant sample, two complementary approaches were used for (2). First, a 95% prediction interval approach treated each infant’s colocalization values as a potential new observation from the adult distribution, following Crawford and Howell [13]. Second, for the pooled infant group (*n* = 2) vs. adults (*n* = 16), Mann-Whitney *U* tests and permutation tests were conducted.

Two-sided Mann-Whitney *U* tests were used for all comparisons. Effect size was calculated for all comparisons as Hedges’ *g* and interpreted as small (|*g*| < 0.5), medium (0.5 ≤ |*g*| < 0.8), or large (|*g*| ≥ 0.8). Permutation tests (10,000 iterations) were performed for all comparisons as validation for Mann-Whitney *U* test. Bonferroni correction was applied within each comparison to the number of variables tested.

Composite niche variables and individual cell-type metrics were also tested as separate families, with Bonferroni corrections applied within each family (6 tests for composites, 13 tests for individual cell types). This approach provided less conservative thresholds than the combined 19-test correction used in the primary analysis.

#### Random forest classifier

To assess whether T-cell colocalization patterns could discriminate younger pre-menopausal females from other adults in a multivariate framework, an exploratory random forest classifier was trained on the 13 individual cell type Pearson correlation values. The classifier used 100 decision trees with a maximum depth of three and a fixed random seed. Infant samples were excluded from classification to restrict the analysis to the adult comparison. Leave-one-out cross-validation was employed to determine younger female classification accuracy, sensitivity, and specificity.

#### Software

Analyses were performed in Python 3.12.3 using pandas (v2.3.3), NumPy (v2.4.0), SciPy (v1.16.3), scikit-learn (v1.8.0), matplotlib (v3.10.8), and seaborn (v0.13.2).

### Immunofluorescence staining

Frozen lumbar DRGs were embedded in optimal cutting temperature compound (Fisher Scientific, 23-730-571), cryosectioned at 20 μm section thickness, and mounted on SuperFrost Plus charged slides (Fisher Scientific, 12-550-15). Each sample included at least two nonconsecutive experimental sections for technical replicates, and each staining run included at least one negative control section which received no primary antibody.

Slides were stored short-term in a -80°C freezer until staining. Slides were transferred to a 37°C laboratory oven to thaw for 1 minute, before immediate submersion into either 10% formalin (human CD3-peripherin, pig, and non-human primate samples) or 4% PFA in 1X PBS (fresh-frozen rat and mouse samples and staining for human residence markers) for 15 minutes. Mouse DRGs that were perfused prior to flash freeze were not post-fixed. All slides were then submerged in an ascending series of ethanol (50%, 70%, and 2 × 100% ethanol) for 5 minutes each. Slides were then air dried at room temperature for 3-5 min, and hydrophobic boundaries were drawn to separate individual sections using a hydrophobic pen (ImmEdge PAP Pen, Vector Laboratories). Slides were placed into a covered humidity control tray. Rat samples were incubated in 10% rat serum (Sigma-Aldrich, R9759) in 1X PBS for 30 min. All samples were then incubated with permeabilization and blocking solution (0.3% Triton X-100 in 1X PBS for fresh frozen mouse DRGs, 10% normal goat serum with 0.3% Triton X-100 in 1X PBS for all other samples) for 1 hour at room temperature. Primary antibody solution was pipetted to cover each experimental section, while negative control sections received blocking solution only. Slides were left covered overnight at 4°C. The primary antibodies and other reagents used are listed in **Supplemental Table 10**.

Following overnight incubation, slides were washed three times with 1X PBS before incubation with secondary reagent solution including DAPI (1:5000, Cayman Chemicals 14285) for 3 hours for residence markers and 1 hour for all other stains. All slides were then washed three times with 1X PBS. Each section, other than those from rat, was then covered with the lipofuscin autofluorescence quencher True Black (Biotium #23014, diluted 1:20 in 70% ethanol) for 1 minute at room temperature. Slides were then washed three times with Milli-Q water, air dried, coverslipped with Prolong Gold antifade mounting medium (Invitrogen, P36930), allowed to cure for 24-48 hours at room temperature, and stored long-term at 4°C.

### Microscopy and T-cell quantification

All slides were imaged under epifluorescence at 20X magnification using a VS200 slide scanning microscope (Evident Scientific). Negative control sections were imaged simultaneously to determine background signal in each channel.

Image analysis was conducted by a researcher blinded to conditions, though true blinding between species was not feasible due to obvious qualitative differences in section size. T-cell quantification was conducted in QuPath (version 0.5.1) [7]. Satellite glial cells were occasionally immunoreactive for CD3, albeit at much lower levels than T-cells. To exclude satellite glia from cell counts in rodent tissues, high and low limits for CD3 channel brightness were set in QuPath according to the optimal conditions for visualizing CD3^hi^ cells in positive control tissue (spleen), which were stained and imaged under identical conditions. Positive control images are presented in **Supplemental Figure 3**.

Peripherin and DAPI staining were used to visualize the neuron-rich region of the DRG bulb, after which the brush tool was used to demarcate an encompassing ROI for each section and measure the region area. T-cells within this region were counted if it displayed a clear halo-like CD3 expression around a singular DAPI-stained nucleus and a diameter of approximately 5-10 μm. When numerous T-cells were aggregated closely together, only distinct and accurately quantifiable T-cells were counted. The points annotation tool was used to generate T-cell counts. At least one section was analyzed per sample. For samples for which multiple sections were analyzed, each was analyzed separately and the counts aggregated (total cell count sum divided by total area of ROIs in mm^2^) to yield total cell density. Representative images were taken by exporting viewer content snapshots from QuPath.

### Statistical analysis

Statistical analysis was conducted in *R* (ver. 4.4.1) in RStudio. Statistical significance was evaluated with Welch’s unequal variance *t*-tests or analyses of variance (ANOVAs), where appropriate, as implemented in the *Rstatix* package (ver. 0.7.2). Statistical significance was evaluated to a level of α = 0.05.

## Notes

### Competing Interest Statement

TJP is co-founder of 4E Therapeutics, PARMedics, NuvoNuro, Nerveli, and Ted and Gregs.

