## Supplemental Figures and Table Legends for "T-cell distribution in the dorsal root ganglion across species, sex, and age"

### Contents

**Supplemental Figure 1.** Predicted T-cell proportion per Visium barcode sorted by species.

**Supplemental Figure 2.** T-cell mean product colocalization scores per dorsal root ganglion cell type compared between human adults and infants.

**Supplemental Figure 3.** Positive control for mouse CD3 immunofluorescence.

--

**Table legends for Supplemental Tables 1-10.**

Supplemental tables are available in separate spreadsheet files.

### Supplemental Figures

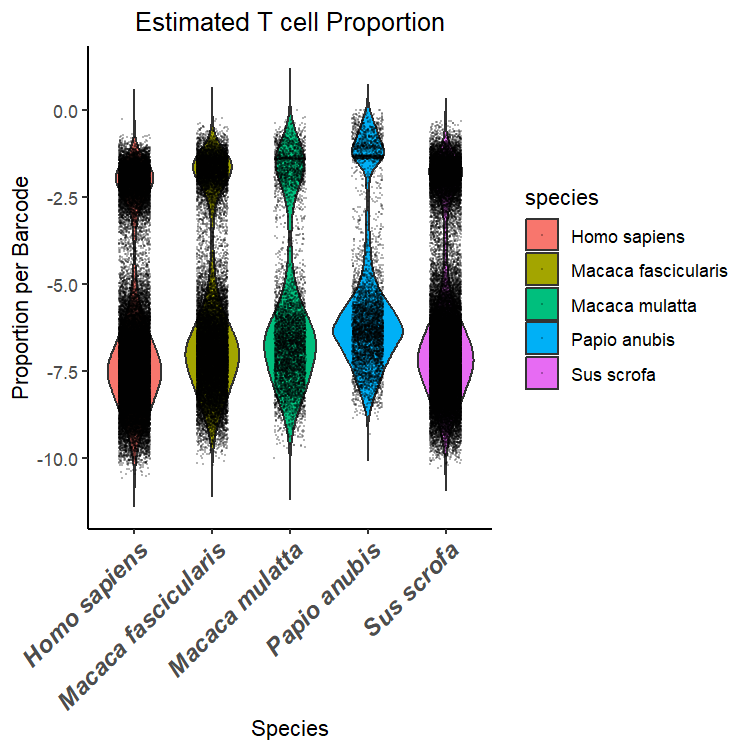

**Supplemental Figure 1.** Predicted T-cell proportion per Visium barcode sorted by species. Each point represents a single barcode from a dorsal root ganglion section from that species, with the y-axis showing the log_10_-transformed predicted proportion. There was a clear bimodal distribution in the estimated T-cell proportion in each barcode for all species at a similar threshold. The threshold of -4 was therefore used to distinguish T-cell ‘positive’ barcodes (>-4) and ‘negative’ barcodes (<-4).

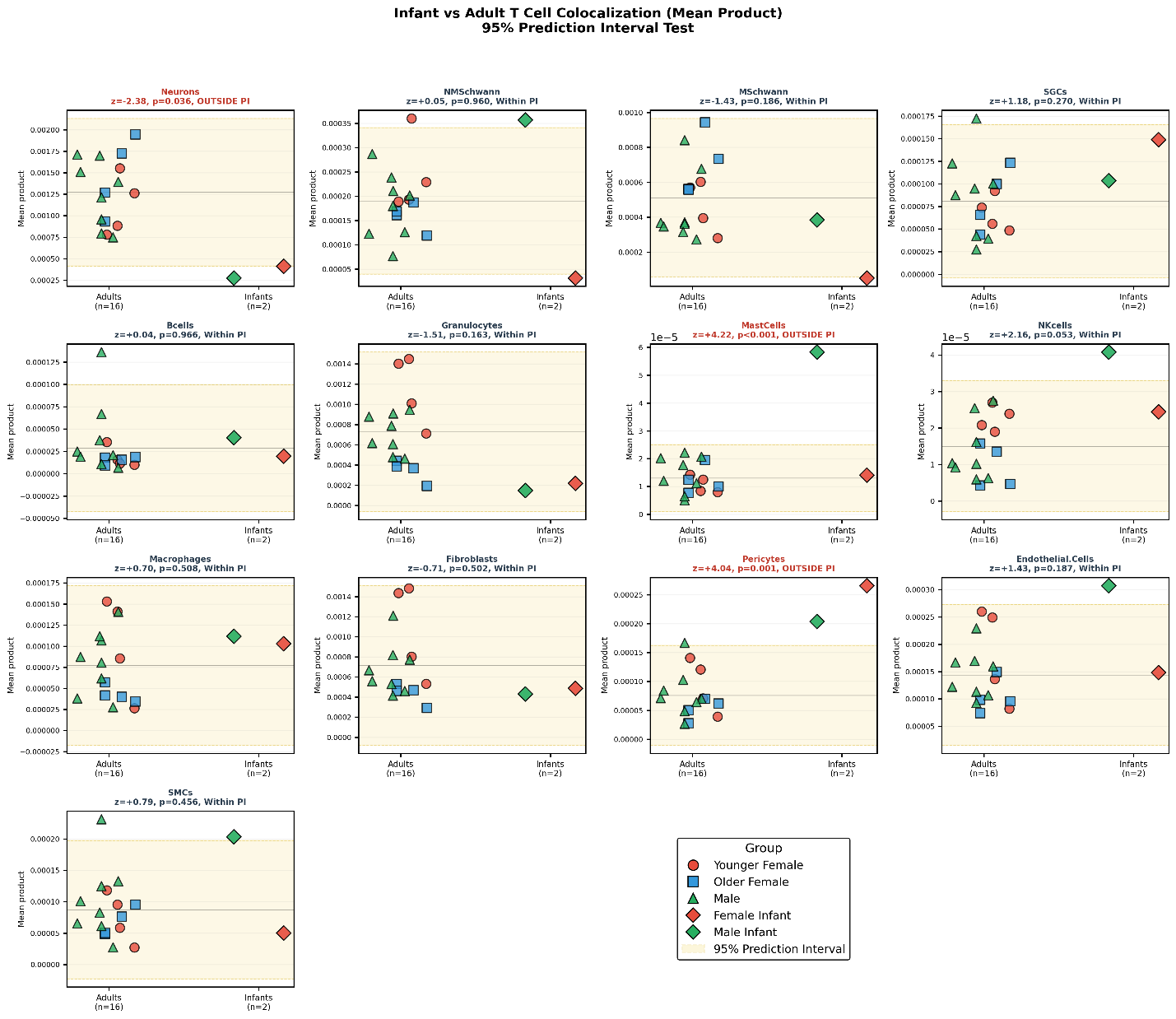

**Supplemental Figure 2.** T-cell mean product colocalization scores per dorsal root ganglion (DRG) cell type compared between human adults and infants. Adults and infants were compared across 13 cell types in the DRG based on the 95% prediction interval (PI) test. T-cell associations in infant DRG significantly below the PI in neurons, and significantly above the PI in mast cells and pericytes. There were no differences in other cell types by this metric.

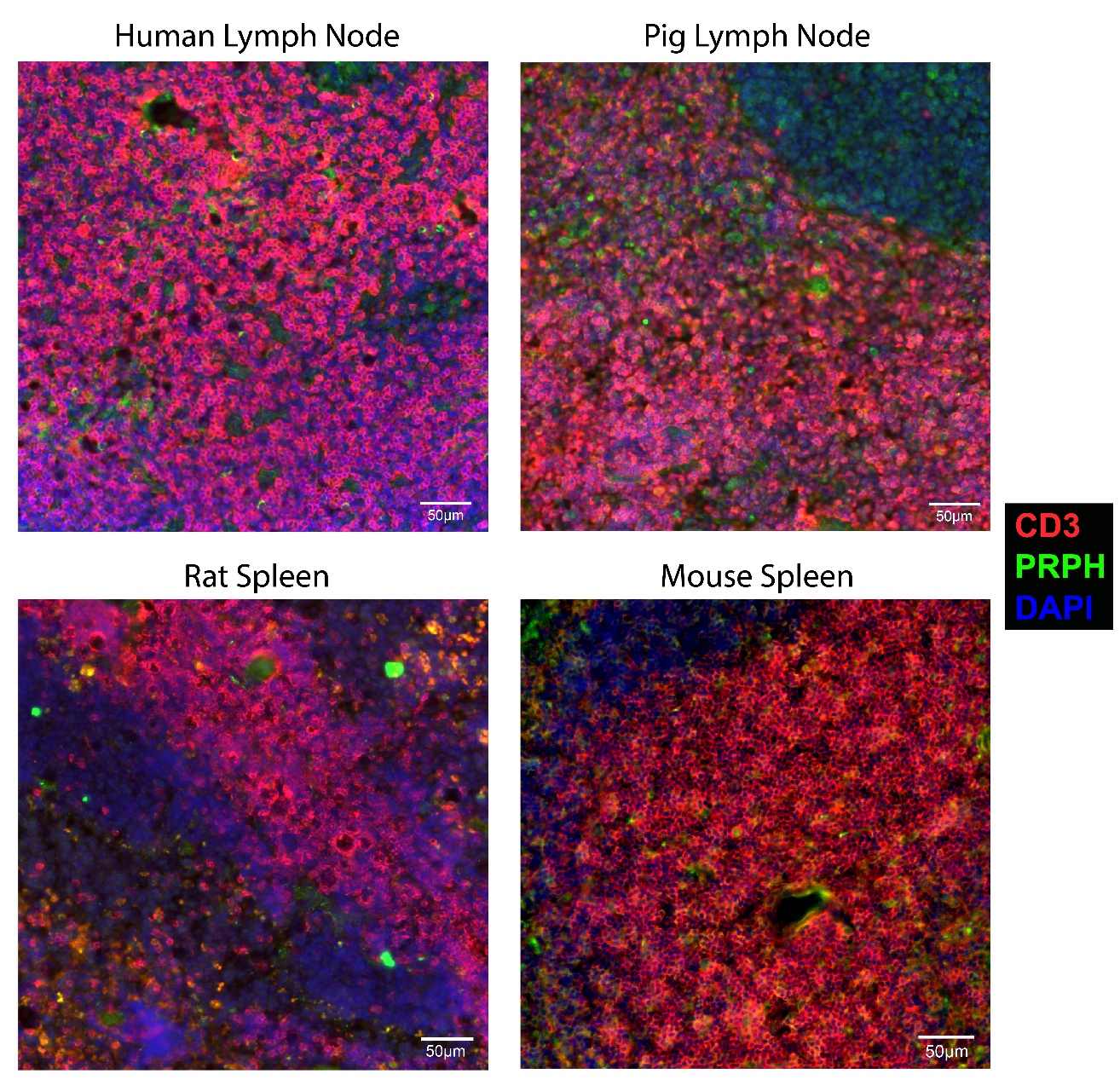

**Supplemental Figure 3.** Positive control images for CD3 immunofluorescence using human lymph node, pig lymph node, rat spleen, and mouse spleen (10X magnification). Tissue was processed identically to DRG sections of the corresponding species.

### Supplemental Table Legends

**Supplemental Table 1.** T-cell-related genes for first-pass manual identification of T-cell-positive barcodes in human Visium data.

**Supplemental Table 2.** Statistical analysis for Pearson correlation scores between T-cells and other human dorsal root ganglion (DRG) cell types. Results for all 19 variables (13 individual cell types and six composites) across all comparisons are presented at both sample and donor levels. Multiple comparison corrections are shown under four schemes: Bonferroni and Benjamini-Hochberg across all 19 variables, and Bonferroni and Benjamini-Hochberg within variable families (6 composites or 13 cell types). Permutation p-values are from 10,000 iterations (seed = 42). Hedges’ *g* uses the standard pooled variance estimator.

**Supplemental Table 3.** Statistical analysis for mean product colocalization scores between T-cells and other human dorsal root ganglion (DRG) cell types. Results for all 19 variables (13 individual cell types and six composites) across all comparisons are presented at both sample and donor levels. Multiple comparison corrections are shown under four schemes: Bonferroni and Benjamini-Hochberg across all 19 variables, and Bonferroni and Benjamini-Hochberg within variable families (6 composites or 13 cell types). Permutation p-values are from 10,000 iterations (seed = 42). Hedges’ *g* uses the standard pooled variance estimator.

**Supplemental Table 4.** Dorsal root ganglion sample details.

**Supplemental Table 5.** Human DRG flow cytometry dissociation times and results.

**Supplemental Table 6.** Flow cytometry primary antibodies.

**Supplemental Table 7.** QC metrics for non-human primate 10x Visium spatial transcriptomic data.

**Supplemental Table 8.** Quality control metrics for human infant 10x Visium spatial transcriptomic data.

**Supplemental Table 9.** Conservation of cell-type marker genes across species. Marker genes from the human single-nucleus RNA-sequencing reference used for SONAR deconvolution were assessed for orthologs in non-human primate (NHP; *Macaca fascicularis*, *Macaca mulatta*, *Papio anubis*) and pig (*Sus scrofa*). Conservation is reported at five thresholds: (1) has pig ortholog, (2) has NHP ortholog, (3) has ortholog in at least one non-human species, (4) has ortholog in both non-human species, and (5) strict conservation (identical gene symbol in human, pig, and NHP). Each cell type has ~250 marker genes from differential expression analysis of the reference dataset.

**Supplemental Table 10.** Immunofluorescence staining reagent details.
